# Prevalence of electricity production among culturable bacteria

**DOI:** 10.64898/2026.07.07.736961

**Authors:** Theo Hembury, Thomas P. Smith, Md Tabish Noori, Klaus Hellgardt, Thomas Bell

## Abstract

Microbial fuel cells (MFCs) technology offers sustainable electricity production. Current research largely focuses on few select model organisms, therefore the true prevalence of exoelectrogenesis amongst bacteria remaining largely unknown. We present a broad-scale survey of monomicrobial electricity production among environmental bacterial isolates inoculated in MFCs, using model organism *Shewanella oneidensis* MR-1 as a benchmark. Of the assessed taxa, 11-22% displayed exoelectrogenic activity, exceeding current predictions and identifying a further three novel exoelectrogenic species. Phylogenetic analysis based on the 16S sequences enabled the evolutionary relationship between isolates to be visualised, revealing that exoelectrogenesis is non-randomly distributed and phylogenetically conserved. Polarisation studies were implemented, revealing that numerous electron transfer mechanism were being utilised to perform exoelectrogenesis. The results of this study imply that bacterial electricity production is more widespread amongst culturable bacteria than previously estimated, with implications for bioprospecting novel exoelectrogens and predicting electrogenic activity in diverse microbial communities.

## Introduction

Microbial Fuel Cell (MFCs) are devices that utilise bacteria to generate an electrical current ^1–4^. MFCs are of huge economic interest as they are a renewable energy source capable of simultaneous wastewater bioremediation ^5–9^. There is an abundance of research on the electricity producing capabilities of bacterial species belonging to the *Geobacter* and *Shewanella* genera in MFCs, including a detailed understanding of the mechanisms in which they perform extracellular electron transfer ^10–12^. Despite the clear economic and sustainability benefits, not all bacteria have the capacity to produce electricity (exoelectrogenesis), so determining the prevalence bacterial electricity production would be useful for identifying new exoelectrogens and advancing this renewable technology.

There are no standard criteria, such as genetic markers, that can be used to define microbial taxa as exoelectrogenic ^13,14^. Furthermore, current identified exoelectrogens are phylogenetically diverse, adding to the difficulty in predicting the prevalence of exoelectrogenesis ^14,15^. Nonetheless, it is estimated that there are approximately 100 taxa that display exoelectrogenic activity, although this prediction is not based on stringent criteria and does not allow an estimate of the proportion of exoelectrogenic taxa within a given community ^16^. There have been attempts to assess the overall abundance of exoelectrogens within an environment, indicating that up to 7.7% of bacterial taxa are exoelectrogenic ^16^. However, such conclusions have been drawn using 16S rRNA sequencing, where exoelectrogenesis from taxa was inferred by matching the identity of known exoelectrogens to the 16S sequences that were surveyed ^16^. Such an approach suffers from enormous uncertainty: unknown exoelectrogens remain unidentified and the absence of genetic markers renders 16S as a poor diagnostic tool ^13,14^. The prevalence of exoelectrogenic activity in any environment is therefore unknown.

Despite the lack of standardised technique for defining exoelectrogens, there are many traits associated with exoelectrogenesis that can be used to assist in determining whether a taxon is exoelectrogenic. Perhaps the most fundamental is the ability to respire anaerobically, for the presence of oxygen would act as a terminal electron acceptor, consequently hindering electrons from reaching the anode ^17–19^. Exoelectrogenesis is a common trait amongst dissimilatory metal reducing taxa as, under anaerobic conditions, they are capable of using exogenous metals as a terminal electron acceptor (TEA) ^20–24^. When inoculated into an MFC, the electrode acts as the TEA, thus producing an electrical current ^25^. Secondly, a majority of known exoelectrogenic taxa are gram-negative, although some gram-positive taxa have also been identified as exoelectrogens, despite their thick peptidoglycan cell wall being thought to prevent extracellular electrogen transfer ^26^.

There are different mechanisms that exoelectrogens may employ to transport electrons to an electrode, with the most common mode of electron transfer being ’direct electron transfer’ (DET), where exoelectrogens with direct, physical contact with the anode transfer electrons ^27^. Alternatively, exoelectrogens may donate electrons to an anode via mediators, either exogenous or endogenously produced, referred to as ‘mediated electron transfer’ (MET) ^28–38^. While this method of electron transfer is not as common as DET, it does not require specific organelles or direct contact with an electrode to produce electricity, so may inadvertently be a more widespread mechanism of electron transfer in MFCs ^39^.

Many of the traits associated with electricity production are not unique to exoelectrogens and are widespread throughout bacterial taxa, with entire phyla being gram-negative and cytochromes (multi-haem outer membrane redox proteins capable of transferring electrons across the outer cell membrane) being widespread, ^40–42^. Furthermore, for a taxon to be deemed exoelectrogenic, not all of the aforementioned traits are required. From this alone, it may be assumed that exoelectrogenesis is more common than currently estimated, which begs the question of how many bacterial taxa are exoelectrogenic ^13,43,44^.

A greater understanding of the frequency of exoelectrogenesis would be significant for a number of reasons. From a biotechnological standpoint, should electricity-producing traits be common and widely dispersed throughout bacteria then there are opportunities to discover novel exoelectrogens that may outperform current model organisms, or possess other desirable traits, such as more versatile growth conditions, that would make them easier to exploit. Additionally, from an ecological perspective, many MFC applications involve large natural communities, like sewage waste, where inoculating and maintaining a single bacterial taxon would be challenging. Ecological theory makes divergent predictions about the amount and stability of ecosystem processes (like electricity production) when traits are widespread in the community compared to when they are rare. If exoelectrogenesis is rare, the focus should be on identifying and tracking keystone taxa, whereas should exoelectrogenesis be common, the emphasis should be on understanding the distribution of electricity production across taxa.

Those taxa that have been reported as exoelectrogenic are phylogenetically diverse, making it difficult to predict the commonness of exoelectrogenesis based on sequencing data alone ^14,15^. This is perhaps most prominent when comparing the two model MFC organisms, *G. sulfurreducens* and *S. oneidensis* MR-1, as they are phylogenetically distantly related, belonging to differing phyla that diverged approximately 2.5 to 2.8 billion years ago ^45–48^. We therefore predicted that exoelectrogenic activity is more widespread and common. To address this hypothesis, we conducted a survey of exoelectrogenic activity using bacteria collected from soil and sediment environments to characterise the commonness of exoelectrogenic activity while using a model exoelectrogenic organism as a benchmark.

## Results

### Prevalence of electricity production

The isolates used throughout are detailed in Table 1 with their electrical data displayed in Figure 1. A t-test was used to compare electricity production between Silwood and Iceland isolates, displaying no statistical significance between the two environments (*p* = 0.56, df = 13.36, *t* = -0.59). An additional t-tests was used to compare each isolates current density measurement against that of the sterile LB control, with *Aeromonas salmonicida*, *Aeromonas popoffii* and *Aeromonas sobria* being statistically significant following a Bonferroni correction for multiple comparisons. When an unadjusted t-test was used a further three taxa, *Cedecea lapagei, Scandinavium goeteborgense* and *Flavobacterium tyrosinilyticum* were statistically significant. The results of the statistical analysis were in part used to determine the bacterial isolates degree of exoelectrogenesis. Isolates displaying statistical significance post Bonferroni correction are classified as ‘Fully’ exoelectrogenic, meanwhile statistical significance prior to a Bonferroni correction are classified as ‘Mildly’ exoelectrogenic, while isolates displaying no statistical significance are ‘Non’ exoelectrogenic.

**Figure 1:**
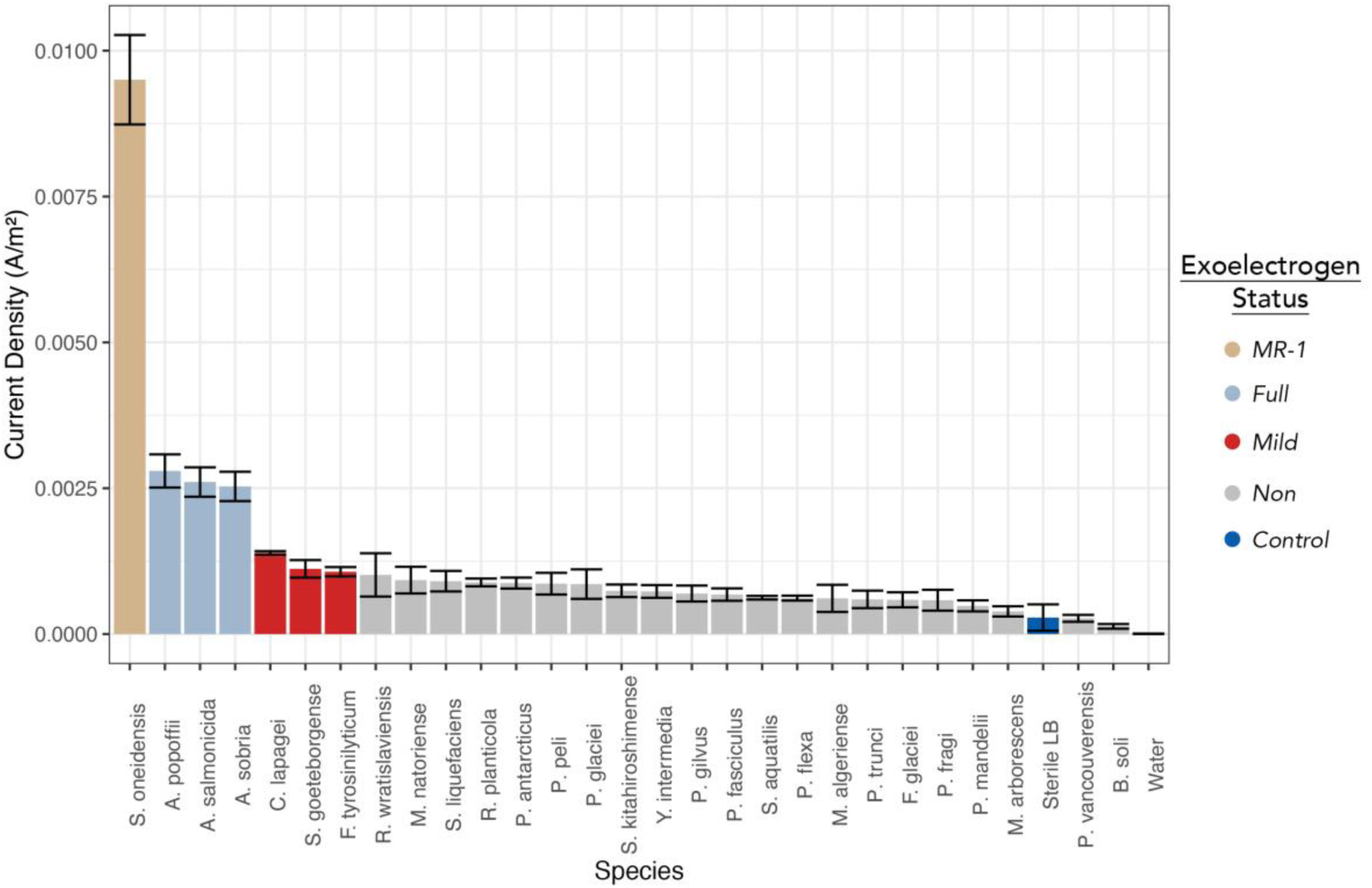
The electrical performance for each of the monocultured isolates with their exoelectrogenic status assigned from the results of the statistical assessment. Current density was calculated using Ohm’s law and electrode surface area measurements. Each replicate for each inoculant was biologically independent, typically being performed in duplicate experiments. A minimum of four independent biological replicates was performed for each isolate with the median current density for each isolate being determined. The displayed data is the mean of these medians with the error bars displaying the standard error of the mean. Further species information is displayed in Table 1.

**Table 1:** List of the isolates assessed and the results of statistical tests comparing current density, as a monoculture, against the current density of the sterile LB negative control. Comparisons were adjusted via the Bonferroni method. (* ≤ 0.05, ** ≤ 0.01, ***≤ 0.001).

| Origin | Taxonomy | Gram Stain | Adjusted P - Value | Df | t |
| --- | --- | --- | --- | --- | --- |
| New York | *Shewanella oneidensis* MR-1 | Negative | 2.3 x 10<sup>-4</sup> *** | 6.99 | 11.53 |
| Iceland | *Aeromonas salmonicida* | Negative | 0.01** | 5.93 | 6.86 |
| Iceland | *Aeromonas popoffii* | Negative | 0.02* | 5.72 | 6.92 |
| Iceland | *Aeromonas sobria* | Negative | 0.02* | 5.94 | 6.64 |
| Iceland | *Pseudomonas peli* | Negative | 1.00 | 5.78 | 1.98 |
| Iceland | *Flavobacterium glaciei* | Negative | 1.00 | 4.77 | 1.17 |
| Iceland | *Phycicola gilvus* | Negative | 1.00 | 4.92 | 1.56 |
| Iceland | *Pseudomonas mandelii* | Negative | 1.00 | 4.02 | 0.82 |
| Iceland | *Pseudomonas vancoverensis* | Negative | 1.00 | 3.41 | -0.06 |
| Iceland | *Serratia aquatilis* | Negative | 1.00 | 3.11 | 1.49 |
| Iceland | *Yersinia intermedia* | Negative | 1.00 | 4.32 | 1.78 |
| Iceland | *Bacillus soli* | Positive | 1.00 | 3.19 | -0.66 |
| Iceland | *Microbacterium arborescens* | Positive | 1.00 | 3.92 | 0.44 |
| Iceland | *Prolinborus fasciculus* | Positive | 1.00 | 4.27 | 1.58 |
| Silwood | *Cedecea lapagei* | Negative | 0.43 | 3.11 | 4.85 |
| Silwood | *Flavobacterium tyrosinilyticum* | Negative | 0.95 | 3.72 | 3.27 |
| Silwood | *Scandinavium goeteborgense* | Negative | 0.73 | 5.22 | 3.07 |
| Silwood | *Raoultella planticola* | Negative | 1.00 | 3.52 | 2.55 |
| Silwood | *Serratia liquefaciens* | Negative | 1.00 | 5.65 | 2.18 |
| Silwood | *Pedobacter trunci* | Negative | 1.00 | 5.20 | 1.15 |
| Silwood | *Psychrobacillus glaciei* | Negative | 1.00 | 5.93 | 1.69 |
| Silwood | *Pseudomonas fragi* | Negative | 1.00 | 5.69 | 1.03 |
| Silwood | *Sphingobacterium kitahiroshimense* | Negative | 1.00 | 4.29 | 1.83 |
| Silwood | *Microbacterium algeriense* | Positive | 1.00 | 6.00 | 1.02 |
| Silwood | *Microbacterium natoriense* | Positive | 1.00 | 6.00 | 2.00 |
| Silwood | *Paeniglutamicibacter antarcticus* | Positive | 1.00 | 4.00 | 2.41 |
| Silwood | *Rhodococcus wratislaviensis* | Positive | 1.00 | 4.96 | 1.68 |
| Silwood | *Priestia flexa* | Variable | 1.00 | 3.21 | 1.45 |

### Phylogenetic signal of electricity production

The phylogenetic tree in Figure 2 shows that novel exoelectrogenic taxa belong to the Pseudomonadota phyla, with the exception of mildly exoelectrogenic taxon, *Flavobacterium tyrosinilyticum,* which belongs to Bacteroidota. These phyla subsequently inform that all novel exoelectrogens are gram negative. From the taxa’s position within the phylogenetic tree, full exoelectrogens are most phylogenetically similar to the *S. oneidensis* MR-1 benchmark. Phylogenetic analysis informs that electricity production was non-randomly distributed across the phylogeny of isolates, determined by testing for phylogenetic signal in the voltages produced by each species using K and λ (see methods). A strong phylogenetic signal was observed using both metrics (λ = 0.95, *p* = 6.59 x 10^-7^; K = 0.57, *p* = 0.005). The results of both the Blomberg’s and Pagel’s method show a strong significance, indicating that exoelectrogenesis is an evolutionary inherited trait.

**Figure 2:**
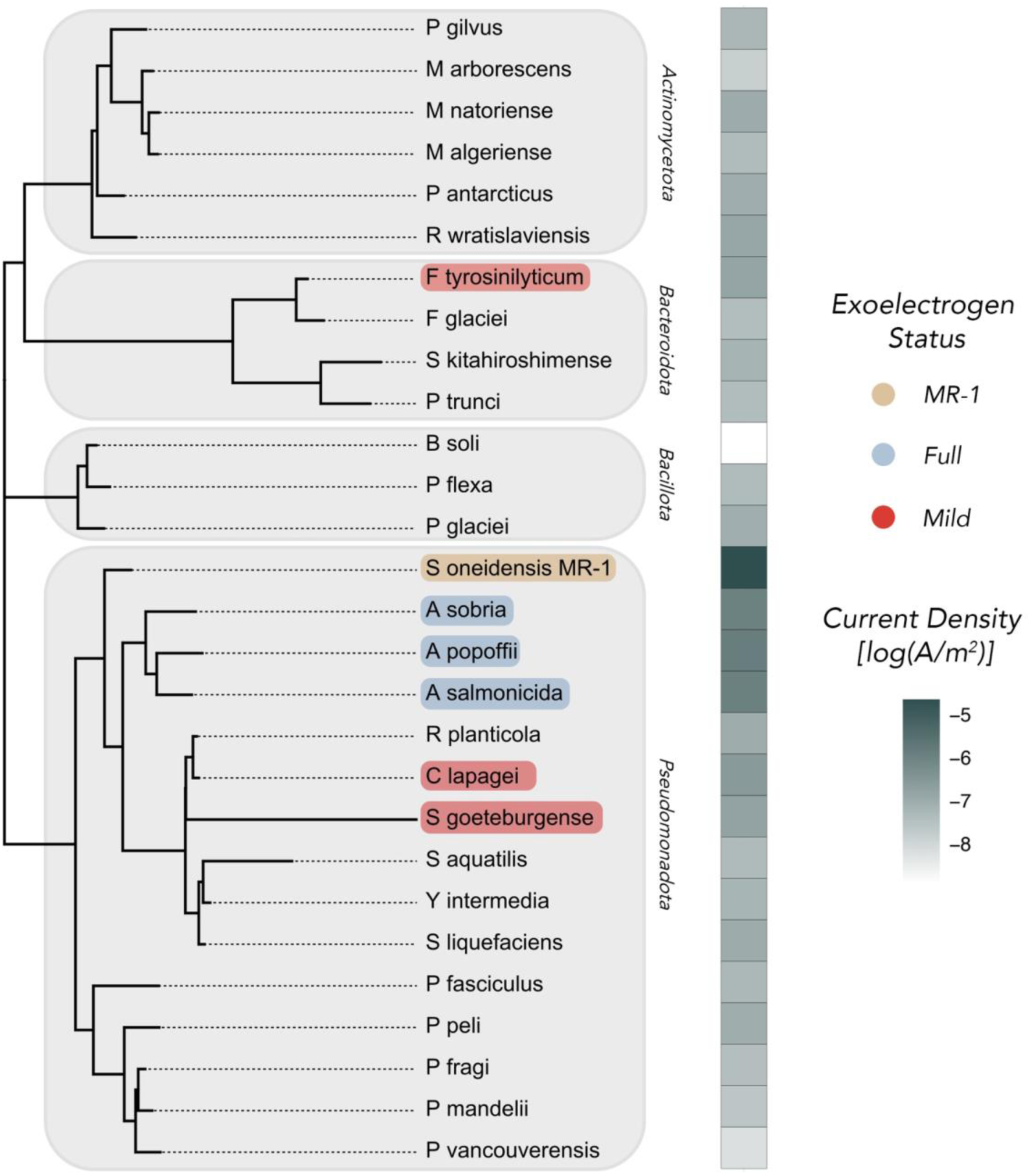
Phylogenetic tree displaying all the isolates used in the experiment, constructed using DNA sequencing of the 16S locus. Light grey boxes surrounding isolates denote the phyla in which those isolates belong. Individual highlighted isolates denote the level of exoelectrogenic classification, concluded from the results of prior statistical testing. Adjacent to the phylogenetic tree is a heatmap displaying the Current Density (from Figure 1), on a log scale, for ease of viewing.

### Efficiency of electricity production

Exoelectrogenic efficiency for novel exoelectrogenic isolates was determined by mapping electricity production per unit of metabolism and biomass, displayed in Figure 3. Mild exoelectrogens maintain their population, denoted as CFU, throughout experimentation while other isolates population declined after the mid-point. Generally speaking, *S. oneidensis* MR-1 has the lowest overall population size whilst *A. salmonicida* has the largest. Despite its low population size, *S. oneidensis* MR-1 maintains a high current density per cell, increasing throughout. Alternatively, all novel exoelectrogenic taxa show little increase in current density on a cellular level.

**Figure 3:**
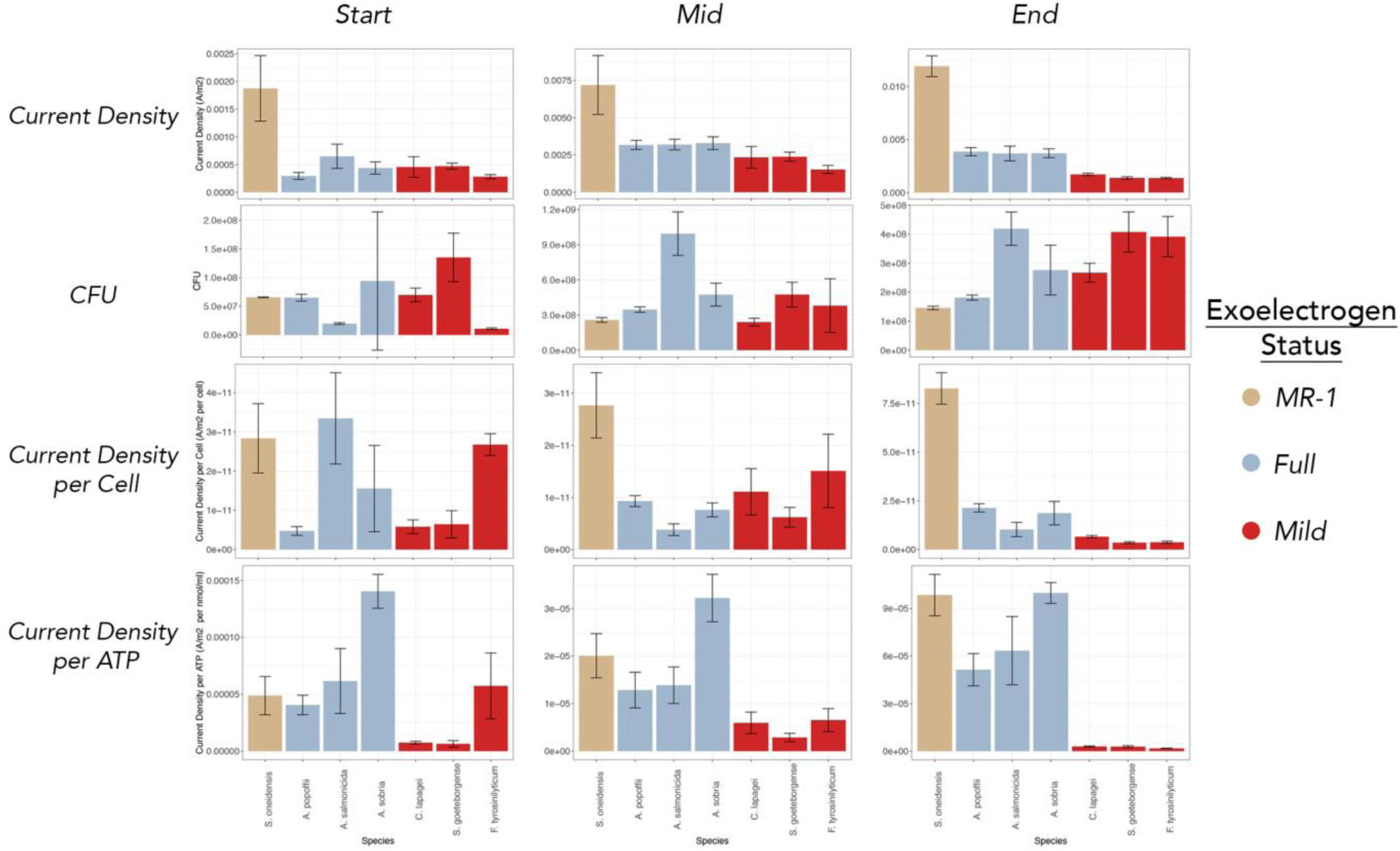
The efficiency of electricity production for select taxa. Current density was calculated using Ohm’s law and electrode surface area measurements described in the methods. The displayed data is the mean from data 1 hour surrounding each time point (see methods) with the error bars displaying the standard error of the mean. The cellular and ATP level calculations were determined used the CFU and ATP measurements, detailed in the methods.

When quantifying current density per ATP, there is an increase throughout experimentation for *S. oneidensis* MR-1. At the start of MFC operation, *S. oneidensis* MR-1 was a neither the worst nor best current producer per ATP however by the end of experimentation is one of the highest performing taxa. An increase in current density per ATP throughout experimentation is also observed with the fully exoelectrogenic isolates. However, the same cannot be said with mild exoelectrogens, as they instead show a decrease towards the end of experimentation, overall performing the worst out of all the assessed taxa.

### Electrical performance of exoelectrogens

Polarisation analysis was conducted to further characterise the electrical performance of the top-performing isolates along with the *S. oneidensis* MR-1 benchmark (Figure 4). *C. lapagei* exhibited the highest ohmic losses, reflected by its steep voltage drop across a narrow current density range (0–∼2.5 mA/m²), followed by *A. popoffii* (∼2.5–5 mA/m²) and *S. oneidensis* MR-1, which sustained the broadest current density range (∼5–9 mA/m²) before collapse. However, mass transfer losses followed the inverse pattern. All three isolates displayed a significant polarisation overshoot, which is visible in Figure 4A as each curve reverses sharply back on itself at low voltages, further confirming that mass transfer limitations were universally high across the assessed taxa. Further details on polarisation curves and their interpretation are in the supplementary materials (Figure 2).

**Figure 4:**
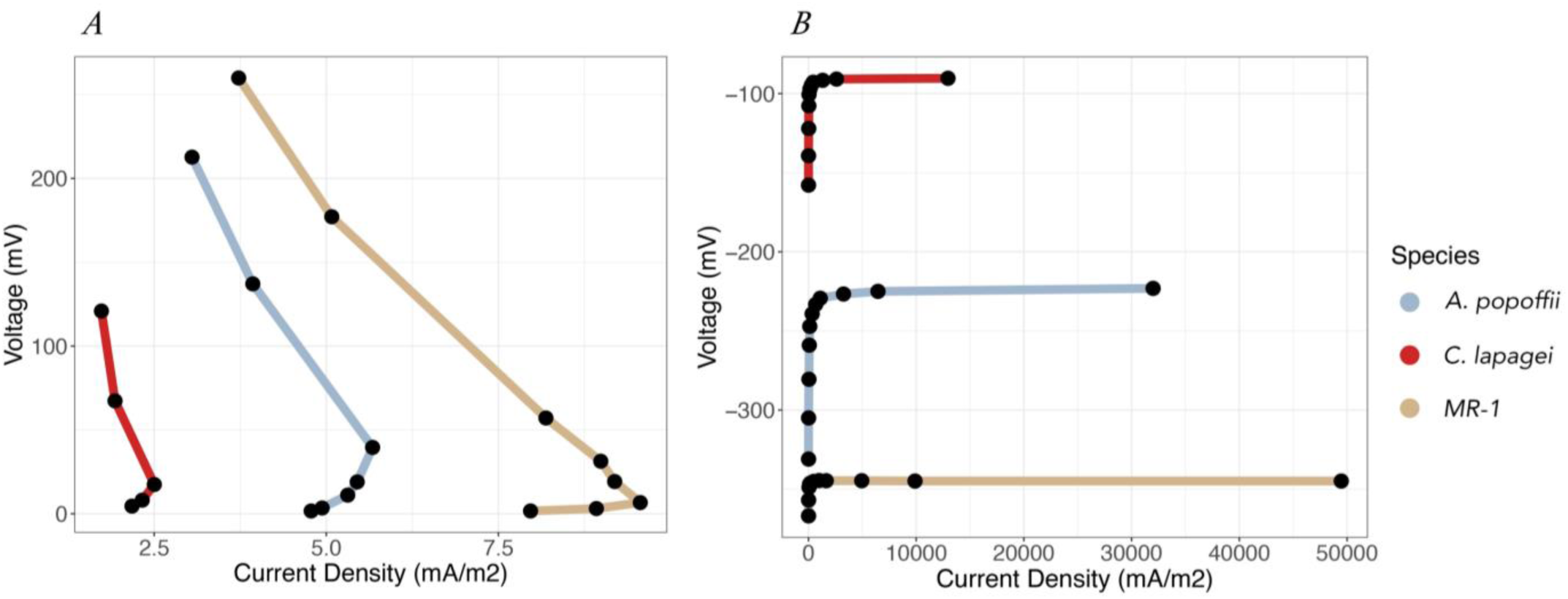
Polarisation curves for the topmost electricity producers from each environment, including the *S. oneidensis* MR-1 benchmark organism. (A) displays whole cell polarisation data whereas (B) displays half-cell polarisation data, formulated by comparing the anode against a Ag/AgCl reference electrode. Note: some polarisation curves in (A) have differing numbers of datapoints. All polarisation curves were collected using the protocol described, however at lower resistances (typically ≤ 50 Ω) current density/voltage would fluctuate and be inaccurate as taxa and/or communities were stressed.

Anodic half-cell potentials (Figure 4B) reveal that *S. oneidensis* MR-1 maintained the flattest and most stable trajectory, spanning the highest current density range (∼50,000 mA/m²) at ∼−345 mV, indicating efficient electron transfer to the anode. *A. popoffii* stabilised at ∼−230 mV across a current density range of ∼30,000 mA/m², while *C. lapagei* maintained a tighter potential around −90 mV but achieved only ∼2,500 mA/m², roughly 8% of *S. oneidensis* MR-1 and 40% of *A. popoffii*, suggesting a physiological constraint on current generation despite anodic stability. Cathode potentials derived from the standard cell potential equation (equation 2) were consistently low across all taxa (Table 2), identifying the cathode as a principal bottleneck for electricity generation.

**Table 2:**
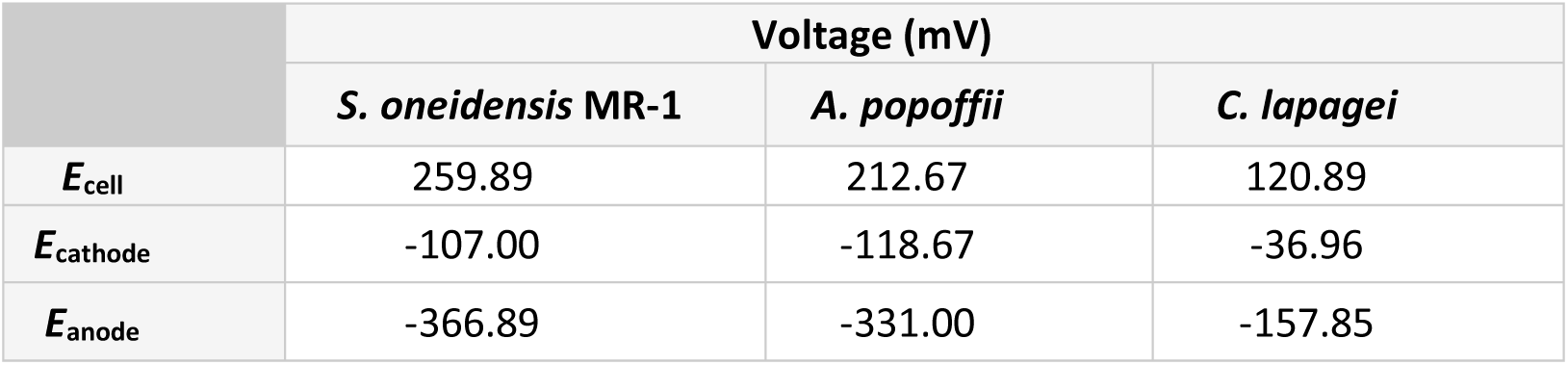
The cell potentials for the differing components of each MFC. The data has been formulated using the data displayed in Figure 4 in conjunction with the standard cell potential equation. All displayed voltages are when MFCs were connected with a 10K Ω resistor. All half-cell potentials are noted against Ag/AgCl reference electrode.

|  | Voltage (mV) |  |  |
| --- | --- | --- | --- |
|  | *S. oneidensis* MR-1 | *A. popoffii* | *C. lapagei* |
| $E_{\text{cell}}$ | 259.89 | 212.67 | 120.89 |
| $E_{\text{cathode}}$ | -107.00 | -118.67 | -36.96 |
| $E_{\text{anode}}$ | -366.89 | -331.00 | -157.85 |

## Discussion

To our knowledge, this is the largest experimental assessment of bacterial isolate electricity production to date. The absence of any definitive criteria for assigning taxa as exoelectrogenic or known genetic indicators means this study sheds light on the prevalence of exoelectrogenic taxa by comparing electricity production against well documented exoelectrogen, *S. oneidensis* MR-1 ^14^. The obtained electrical data displays that nearly all taxa had a detectable current density, as predicted, due to microbial redox molecules ^13^. None of the assessed taxa displayed a current density greater than that of *S. oneidensis* MR-1. Of the assessed isolates, three taxa, all of which are Aeromonas spp, were statistically significant electricity producers following adjustment for multiple comparison and therefore classified as being fully exoelectrogenic. To our knowledge this is the first recorded case of full exoelectrogenesis from *A. popoffii, A. salmonicida and A. sobria,* as the only other recorded exoelectrogen from this genus is *Aeromonas hydrophila* ^49^. A further three isolates, all originating from Silwood, were classified as being mildly exoelectrogenic due to them displaying statistical significance, compared to the LB control, prior to a Bonferroni correction.

The three novel, fully exoelectrogens alone represent 11% of the total assessed isolates (excluding the *S. oneidensis* MR-1 benchmark), surpassing previous estimates which suggest fewer than 8% of taxa are exoelectrogenic when assessing varying soil environments ^16^. Furthermore, estimates suggest there are approximately 100 known exoelectrogenic taxa ^13^. Relative to the findings from prior studies, the increased exoelectrogenic prevalence displayed in this study suggests that current estimates are lower bound, as electricity production is more common than initially believed, although still not a widespread trait. Irrespective of the percentage of species that are exoelectrogenic, identifying taxa that can produce current was a straightforward task which may be testament to the trait’s prevalence. We accept that assessing two environments is not representative of all Earth’s biomes, although assessing two environments broadened the diversity of analysed taxa. Determining which environment is host to the most and best exoelectrogenic taxa is outside the remit of this study.

There is no significant difference between the two environments that were assessed, indicating that the presence of novel exoelectrogens is independent of the environment in which they reside. This outcome is not anticipated, as exoelectrogenesis is an anaerobically driven process, from which it may be inferred that anaerobic environments, such as the sediment in which the Icelandic taxa were isolated, would-be home to more exoelectrogenic taxa ^19^. Despite the lack of significant difference between the two assessed environments, there is variation in exoelectrogenic capability between environments, as the Icelandic taxa have been assigned as fully while the Silwood taxa as mildly exoelectrogenic, supporting the assumption that anaerobic environments would support exoelectrogens.

For the most part, all of the novel exoelectrogenic taxa are in close proximity to the *S. oneidensis* MR-1 on the phylogenetic tree displayed in Figure 2, with fully exoelectrogenic taxa in closer proximity than those classified as mild. This suggests that there is a relatively common ancestry for exoelectrogenic isolates which is supported by the results from both a Blomberg’s and Pagel’s phylogenetic analysis, informing that electricity production amongst taxa is an inherited trait. This result is noteworthy for it highlights that in bacterial electricity production is a conserved trait, which is a step in the right direction in discovering methods of predicting exoelectrogenesis amongst taxa.

Mildly exoelectrogenic taxa are more phylogenetically diverse than those that are fully exoelectrogenic as they span multiple genera and phyla. However, one thing that all the novel exoelectrogens have in common is that they are all gram-negative, which is a common, but not fundamental, trait amongst exoelectrogens ^50^. With the exception of *F. tyrosinilyticum*, all novel exoelectrogens are Pseudomonadota, which has been cited as being the phyla in which most exoelectrogens belong ^50^. *F. tyrosinilyticum,* belongs to the phyla Bacteroidota, which has additionally been noted for its abundance and efficient electricity production in MFCs ^51^. The phylogenetic analysis further legitimises the identification of novel exoelectrogens in this study.

A different approach was taken to examine the efficiency of electricity production from each taxa, for the complexity of the LB substrate used in this study raised concerns over the reliability of implementing more conventional coulombic efficiency as a measurement for each bacterial isolate. Nonetheless, this was calculated for each isolate and is displayed in the supplementary materials. The benchmark organism for this study, *S. oneidensis* MR-1, displays a relatively low population size throughout experimentation, yet produces the highest current density out of all the assessed isolates. The high current density on both a cellular and metabolic level indicates that each cell contributes greatly to electrical current and is therefore not reliant on a large population size to produce large current densities. *S. oneidensis* MR-1’s exoelectrogenic efficiency may be attributed to its ability to donate electrons directly and via mediators to an electrode, eliminating any spatial constraints ^30,52,53^. Despite it’s exoelectrogenic efficiency, it takes the length of the experiment for *S. oneidensis* MR-1 to produce a high current density per ATP, suggesting slow and inefficient resource use, in which *S. oneidensis* MR-1 has been previously cited for its inability to metabolise certain substrates ^54–61^.

Broadly speaking, the overall population size (CFU) for fully and mildly exoelectrogenic taxa follow the same trends throughout the time series and, for the most part, exceed that of *S. oneidensis* MR-1. Alternatively, mapping current density to ATP reveals full exoelectrogens are consistently more efficient compared to mild exoelectrogens, albeit, with the exception of *A. sobria*, not as efficient as *S. oneidensis* MR-1. The low current density per cell and per unit of ATP exhibited by mild exoelectrogens suggests less efficient extracellular electron transfer (EET) pathways are being utilised, which further supports their exoelectrogenic classification. In comparison, fully exoelectrogenic taxa have a far greater current density per ATP and overall current density, suggesting that substrate consumption and metabolic activity translate to electrical current more in comparison to the mild exoelectrogens where it translates more into growth. With the substrate being the main carbon source for MFC microbial populations, the choice of substrate will impact the electrical performance ^8^. This greater substrate utilisation from the full exoelectrogens is likely responsible for the differing exoelectrogenic classification.

The polarisation data corroborate the exoelectrogenic classifications assigned to the assessed isolates. For instance, *C. lapagei*, classified as a mild exoelectrogen, displayed a standard anode potential (Table 2) more positive than the thermodynamic limit for efficient exoelectrogenesis (−200 mV), implying reduced electrode colonisation ^28^. Should this reduced colonisation result in MET being the main EET mechanism, it remains unclear whether the electron shuttles facilitating MET are endogenously synthesised or derived from the growth substrate, for the flavin constituents of LB media have been proposed as likely mediators, which would introduce uncertainty regarding *C. lapagei*’s intrinsic exoelectrogenic capacity when cultured under alternative media conditions ^14,62^.

The comparatively high ohmic losses observed in *C. lapagei* are consistent with the additional resistive burden imposed by MET mechanisms and further justify its mild exoelectrogenic designation. In contrast, *A. popoffii*, classified as a full exoelectrogen, exhibited an anode potential with a greater reducing power (more negative) than the thermodynamic limit for efficient exoelectrogenesis (−200 mV), therefore supporting the use of DET for current generation ^28^. This is further corroborated by reports of nanowire structures across *Aeromonas* spp., providing a physical basis for direct electrode contact, although the specific membrane proteins involved remain uncharacterised ^49,63,64^. The combination of nanowire prevalence across the genus and the novel exoelectrogenic *Aeromonas* spp. identified in this study raises the intriguing possibility that electricity-generating capacity may be broadly distributed throughout *Aeromonas*.

## Conclusion

The performance of a broadscale experiment assessing electrical activity amongst diverse taxa has highlighted that the prevalence of exoelectrogenic activity is greater than initially believed, as between 11 and 22% of assessed taxa displayed exoelectrogenesis, all of which being gram negative which aligns with other studies ^50^. None of the assessed isolates produced an electrical current greater than the benchmark organism, *S. oneidensis* MR-1 which displayed efficient exoelectrogenesis, independent of its population size. Statistical significance was used in conjunction with other performance parameters to determine whether isolates were fully of mildly exoelectrogenic. Novel isolates that displayed full exoelectrogenesis are closer related to the *S. oneidensis* MR-1 benchmark than the mildly exoelectrogenic taxa. The use of further phylogenetic tools indicates that exoelectrogenesis is an inheritable trait, therefore showing promise for the identification of future exoelectrogens.

## Materials and Methods

### Bacterial Isolates

We assessed the electrogenic activity of bacterial isolates collected from two locations: anaerobic sediment from a freshwater stream in Iceland, and topsoil from Nash’s Field, Silwood Park (UK) ^65,66^. Both sets of isolates had been identified (16S rRNA locus) prior to this experiment. Details of the isolates are given in Table 1. *Shewanella oneidensis* MR-1 (first isolated from lake sediment taken from Lake Oneida, New York State, USA) has been studied extensively as a model exoelectrogen and so was used as the benchmark for electricity production ^22,50^. Prior to inoculation in MFCs, bacterial isolates were grown at room temperature (20°C ± 2°C) on a shaking platform for at least 72 hours to ensure transition into stationary growth phase. After MFC assembly, 1.5 ml of bacterial culture (1% of the carrying capacity) was inoculated into 148.5ml Luria Broth (LB) assembled in house (Tryptone - Sigma, NaCl - Sigma, Yeast Extract- BioServ). Two negative controls were used: sterile LB and sterile Milli-Q water.

### MFC assembly and data collection

We used two-chamber, glass ‘H-type’ MFCs (Figure 5). The electrodes were carbon brushes with a surface area of 69.73 cm^2^ and a titanium wire core, spaced approximately 112 mm apart and connected via a 1000 Ω resistor. Details of how the surface area was calculated can be found in the supplementary materials.

**Figure 5:**
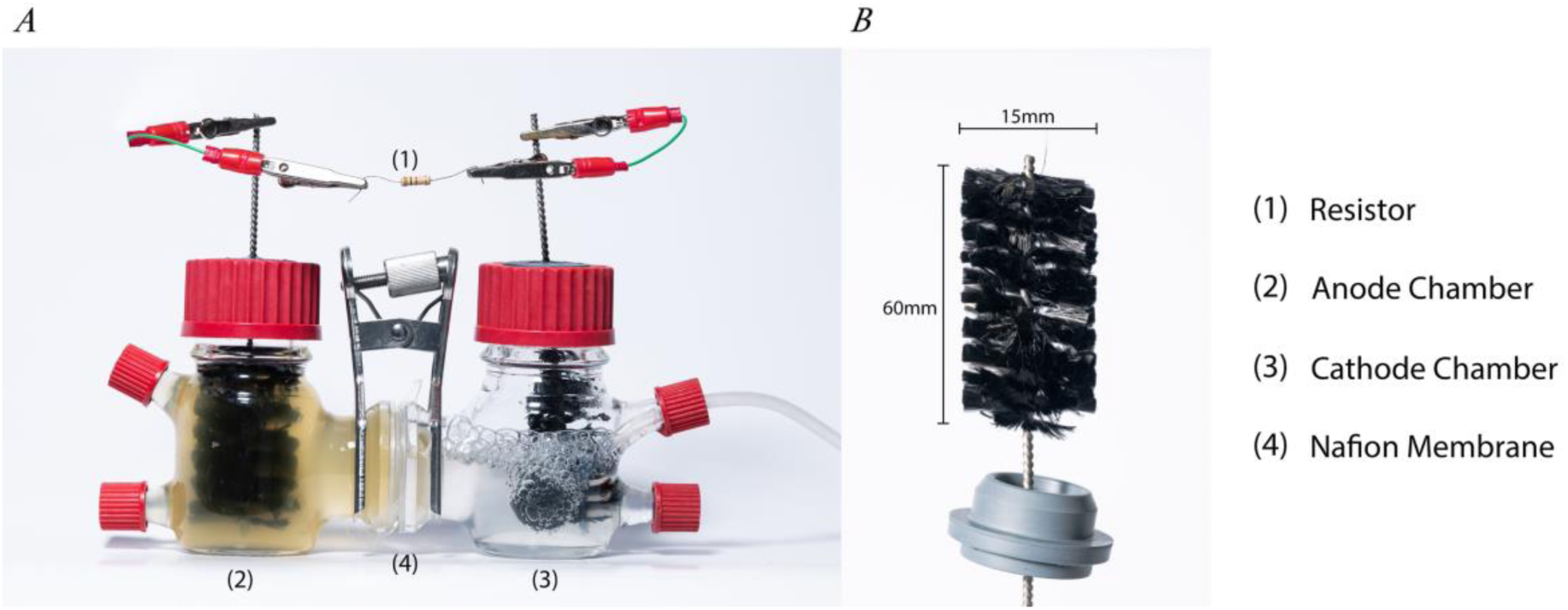
Diagram of the two chamber ‘H-type’ MFC used in this study. (A) Fuel cell with the different key components highlighted. (1) The resistor (2) the anode (3) the cathode (4) Nafion membrane. (B) The carbon brush electrodes used at both the anode and cathode, held in place via a pierced rubber bung.

The Proton Exchange Membrane (PEM) separating the MFC chambers was Nafion 117 (DuPont), which was sterilised using 10% bleach (NaClO) and left to soak for at least 45 minutes prior to rinsing with sterile Milli-Q water. The surface area of the Nafion membrane was approximately 16.65 mm^2^. All other MFC components were sterilised via autoclave and aseptic techniques used when assembling MFCs.

LB growth media was used throughout experimentation due to the wide latitude for growth. To reduce residual oxygen at the anode, chambers were filled with large volumes of media to minimise the headspace in each MFC. Oxygen was used at the cathode, achieved by aerating 100 ml of sterile Milli-Q water with a flow rate of 3.2 L/min. To encourage aeration an air stone was attached to the hose and placed in the cathode chamber. Cathode chambers were topped up as required to maintain volume, typically once every 2 to 3 days.

Both anode and cathode chambers had two sampling ports, closed via screw cap holding a rubber seal to prevent gas exchange. The cathode port was used for aeration while the anode port was used for sampling. MFCs were sampled using a sterile syringe and needle punctured through a rubber port, sterilised with 70% ethanol before insertion. Syringes were pumped and purged to agitate the MFC community.

Voltage was recorded over 164-hours, with the data converted to current using Ohm’s Law, displayed in equation (1), (V = voltage, I = current, and R = resistance) and further converted to current density using the surface area measurements of the electrode. The median voltage across the entire 164-hour time series for each replicate was used as a measure of average electricity production for each MFC. The median was used due to inoculants fluctuating electrical performance during biofilm maturation, as displayed in supplementary materials (Figure 1). The distributions of these voltages are skewed, rendering a mean average inappropriate. Some of these perturbations coincided with sampling and water replenishment at the cathode. A minimum of four biological replicates were performed for each of the assessed isolates, typically being performed in duplicate experiments.

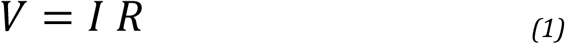

### Bacterial growth and activity

MFCs were sampled at the Start, Middle and End (hours 0, 91 and 164) of experimentation. At each sampling point absorbance and metabolic activity were measured. The absorbance was determined by recording the optical density of 100 μl anodic media at a wavelength of 600 nm (OD_600_). For each timepoint the current density was determined from the voltage data collected one hour surrounding each sampling timepoint.

An aliquot from the liquid monocultures for each species was taken and serially diluted before spreading onto solid LB agar. Simultaneously, aliquots had their absorbance (OD_600_) taken. Solid cultures were left to incubate at room temperature (20°C ± 2°C) for a minimum of three days. After this incubation period, CFU’s were calculated and an average across replicates determined. Using the CFU data coupled with absorbance, a linear model was constructed and used to determine the population size for each isolate at different timepoints by coupling with the experimental absorbance data.

Metabolic activity was measured by quantifying adenosine triphosphate (ATP) in the MFCs using BacTiter-GloTM kit (Promega, Madison, Wisconsin, USA). Further details of this can be found in the supplementary materials.

### Phylogenetic analysis

Following the methods in Smith *et al*., (2022), sequences were aligned to the SILVA 16S database using SINA ^67^. Phylogenies were produced using the 16S sequences of the isolates with RAxML using a general time reversible substitution model with gamma distributed rates. To test evolutionary constraints on the current density produced by each species, two metrics of phylogenetic signal where calculated – Blomberg’s K and Pagel’s λ ^68,69^. Tests for these metrics were implemented in the “phytools” R package. Values of 0 imply no phylogenetic signal, whereas values of 1 imply strong phylogenetic signal (i.e. the distribution of voltages at the tips of the phylogeny are strongly driven by shared evolutionary history). λ is bounded between 0 and 1, with intermediate values showing some evidence for phylogenetic signal. K is not bounded, and values greater than 1 indicate more phylogenetic signal than expected under a null Brownian motion model, implying very strong conservation of traits.

### Polarisation

Full-cell polarisation curves were created for the *S. oneidensis* MR-1 benchmark and the topmost electricity producers from each assessed environment. Monomicrobial cultures were grown for a minimum of 164-hours under the same conditions as those used for the main experiment. Following this, the resistor (1000 Ω) was removed and the MFC left for a further 3 to 4 hours to operate at open circuit voltage until voltage stabilised. Polarisation studies were performed using a variable resistance box. The curves were generated galvanostatically and from three biological replicates, starting at 10 kΩ and progressively decreasing the resistance to 1 Ω, allowing the cell voltage to stabilize at each step before data collection. At each resistance, three voltages were recorded using a multimeter and later converted to the corresponding current and power density using Ohm’s Law. Polarisation curves for isolates were formulated from three biological replicates. The half-cell potentials (anode and the cathode) were measured with respect to a sterilised Ag/AgCl reference electrode and the standard cell potential equation (2).

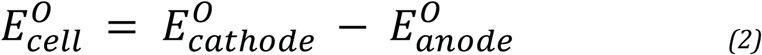

## Acknowledgements

I would like to express my gratitude and thanks to the co-authors in this study: Dr Thomas P Smith, Dr Md Tabish Noori, Professor Klaus Hellgardt and Professor Thomas Bell. I would like thank Imperial College London – Silwood Park for providing me with a wonderful and inspiring place to study and conduct my research.

## Data availability Statement

Data associated with this study is part of this manuscript. Materials associated with this study can be obtained from the corresponding author upon a reasonable request.

