## Supplementary Materials for "Prevalence of electricity production among culturable bacteria"

#### Bacterial Isolates

The Icelandic species were isolated from freshwater stream anaerobic sediment across numerous pristine sites in the Hengill region [1]. The Silwood isolates were obtained from the top 20 cm of soil from Nash's Field long term experiment at Silwood Park, UK [2]. *S. oneidensis* MR-1, was the benchmark species used throughout, first inoculated in lake sediment from Lake Oneida, New York State, USA [3]. BLAST was used to provide species identities based on the 16S sequences for each environmental isolate. The sequencing methods are the same as described in Smith *et al.,* (2024) [1]. The 16S sequence for *S. oneidensis* MR-1 was obtained from the NCBI genbank database (accession number NR_074798).

#### MFC Architecture and Electrodes

The carbon brush electrode surface area was determined to standardise the MFC electrical performance and aid cross-study comparisons. The thickness of individual carbon fibres was 7 µm which is consistent with published studies that report a range of 5-10 µm [4]. The length of the brush was 60 mm, with the carbon fibres extending 14 mm from the titanium wire core. From the thickness and length of each individual fibre, the surface area per fibre was calculated as 0.31 $mm^2$ assuming a cylindrical shape. The total fibre count was determined using light microscopy to extract 100 individual carbon fibres from the electrode prior to being weighed on a precision balance to obtain a weight per fibre. All fibres were then removed from the brush to obtain the total weight of all the fibres and therefore the number of fibres per brush. Adopting this approach, it was noted that each carbon brush electrode comprised of approximately 224, 000 individual fibres and therefore a total surface area of 69.73 $cm^2$. This estimate of surface area per brush is used throughout. It is worth noting that this approach does not take into consideration any irregularities on the surface of the fibres at a microscopic level, so the surface area estimate should be taken as a lower bound.

#### Sterilisation

Aseptic techniques were used throughout as and when required. All autoclaving was performed at 121°C at 2.4 bar pressure for 15 minutes. All water used throughout was Milli-Q water that had been autoclaved prior to use. Apparatus not specified was purchased pre-sterilised and/or assembled aseptically. Port lids and the rubber O-rings, located at the flange of the MFC were assembled and wrapped in foil prior to autoclaving.

Electrodes were placed in autoclave bags for sterilisation via autoclave with anodes and cathodes separated for sterilisation. Electrodes were placed into the bag with the same orientation to prevent accidental contact with the electrode fibres. Air stones at the cathode had their external surface sterilised with ethanol (70%). Hoses were sterilised with ethanol and the interior with sterile water.

Nafion was left to soak in 10% bleach (NaClO) for a minimum of 45 minutes. Following this, under aseptic conditions, bleached membranes were soaked in sterile Milli-Q water. Whilst soaking, the membranes were placed on an orbital shaker to assist in removing bleach residues that may have accumulated on the membrane for a minimum of 45 minutes. This was an important step in Nafion sterilisation as $Cl^-$ ions in an MFC may hinder oxygen reduction at the cathode [5,6].

Sterile (autoclaved) forceps were used to place the Nafion into the MFC. Additional care and precision was taken to grasp the membrane by the outermost edge which would not be in contact with the MFC bacterial community and instead be on the peripheral of the MFC. Forceps were changed frequently between MFC assembly, with a single pair typically being used no more than four times before being substituted. Between use, forcep tips were left submerged in 10% bleach (NaClO). Any containers used, for example those used as a water bath or bleach wash, were autoclaved before use.

#### Data Collection

Current density was determined from the electrode surface area measurements, with Ohm’s law being used to convert the collected voltage data into electrical current. Throughout the 164-hour time series, the median voltage per replicate was used as an alternative to the mean average. This was due to the sharp data fluctuations, displayed in Figure 1.

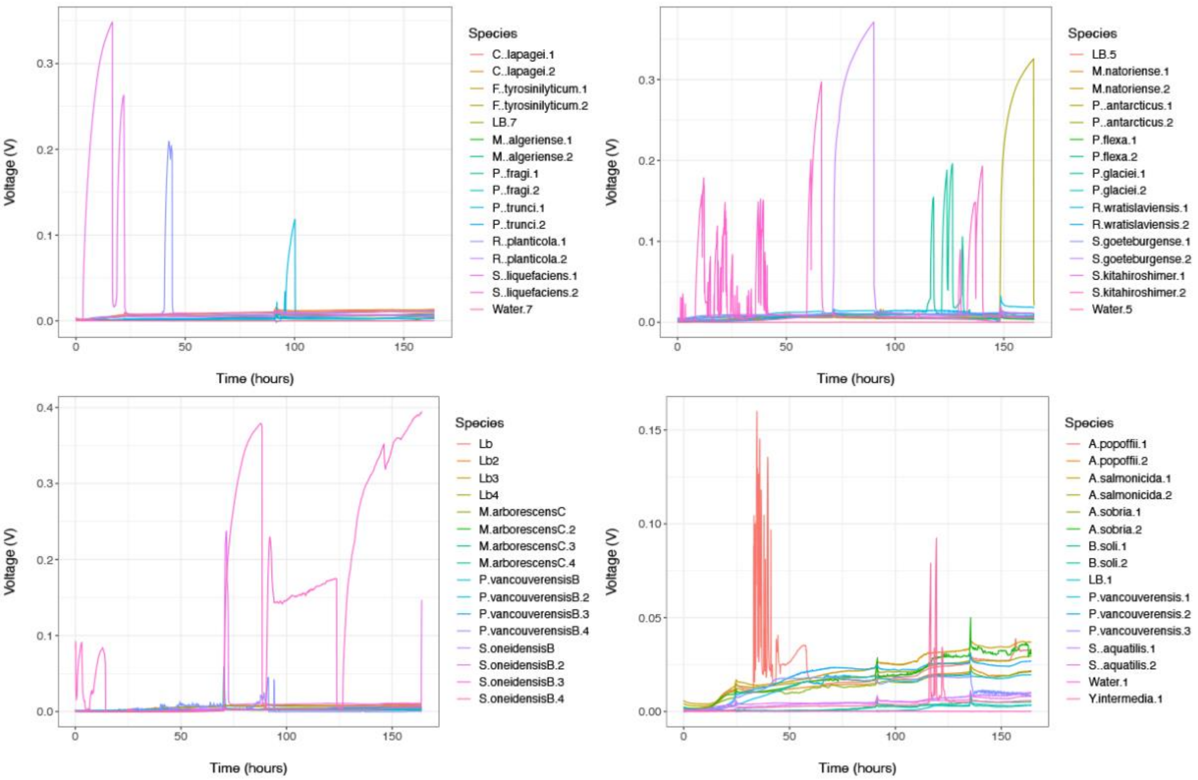


**Figure 1:** The continuous voltage data for selected experiments. The data displayed was formulated using the raw voltage collected in 5-minute intervals. These selected experiments display extreme voltage fluctuations which we negated by displaying the electrical data as a median rather than a mean.

### ATP Assay

ATP was quantified on a 100 µl 1:50 dilution in sterile Milli-Q water. A BioTekTM SynergyTM 2 was used to dispense 50 µl of reagent into the samples which were shaken for five seconds prior to luminescence being measured, five times with 75 second intervals between recordings. To determine the ATP concentration, in nmol/ml, the maximum luminescence reading was taken, and the control luminesce subtracted before dividing by a constant 1693.1 and adjusting for a dilution factor. This is outlined in equation (1) detailed in Mombrikotb *et al.,* (2022) [7].

$$\text{ATP (nmol ml}^{-1}\text{)} = \left(\frac{\text{luminescence}_{\ max} - \text{luminescence}_{\ control}}{1693.1}\right) \text{X dilution factor} \tag{1}$$

#### Polarisation

The implementation of polarisation curves allows the relationship between current (and current density) with voltage to be observed. Supplementary materials Figure 2 displays the distinct regions within polarisation data. Activation losses denote the losses that arise due to the activation energy required for the reaction to take place. Ohmic losses occur due to resistance at different locations of the MFC, for example, the substrate. Mass transfer losses arise due to limitation in transferring substrate away, or to, an electrode.

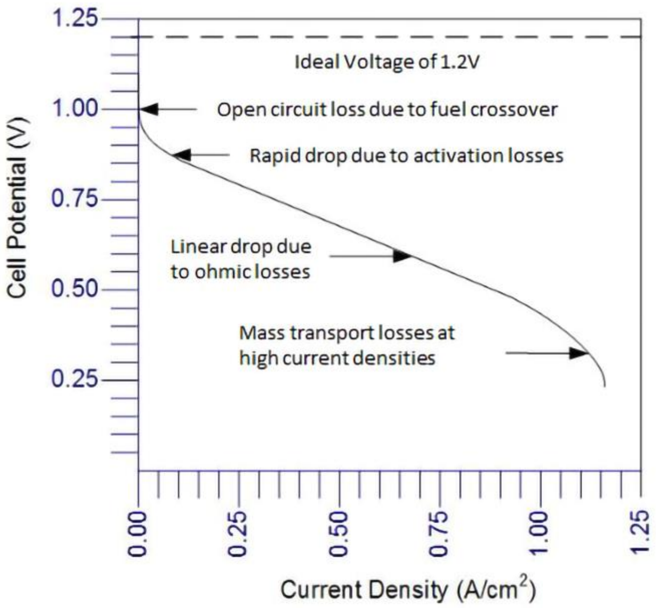


**Figure 2:** A fictitious example of an ideal, typical polarisation curve [8].

#### Statistically Significant Exoelectrogens

The results of statistical analysis aided in conferring whether a taxa is exoelectrogenic. T-tests were used to compare each taxon's current density against that of the LB control, with the results both before (table 1) and after Bonferroni correction used to distinguish between fully and mildly exoelectrogenic taxa. While environment was not statistically significant for electricity production, Figure 3 shows the Current Density for each assessed isolate, however also distinguishes taxa based on their origin.

**Table 1.** The isolates used, inoculated as monocultures into MFCs and their unadjusted p-values. Current Density was compared against the current density of the sterile LB negative control. (* ≤ 0.05, ** ≤ 0.01, ***≤ 0.001).

| Origin | Taxonomy | *p* – value |
|---|---|---|
| New York | *Shewanella oneidensis* MR-1 | $8.41 \times 10^{-6}$ *** |
| Iceland | *Aeromonas popoffii* | $5.55 \times 10^{-4}$ *** |
| Iceland | *Aeromonas salmonicida* | $4.95 \times 10^{-4}$ *** |
| Iceland | *Aeromonas sobria* | $5.87 \times 10^{-4}$ *** |
| Iceland | *Pseudomonas peli* | $9.65 \times 10^{-2}$ |
| Iceland | *Flavobacterium glaciei* | $2.97 \times 10^{-1}$ |
| Iceland | *Phyicola gilvus* | $1.80 \times 10^{-1}$ |
| Iceland | *Pseudomonas mandelii* | $4.59 \times 10^{-1}$ |
| Iceland | *Pseudomonas vancouverensis* | $9.58 \times 10^{-1}$ |
| Iceland | *Serrattia aquatilis* | $2.30 \times 10^{-1}$ |
| Iceland | *Yersinia intermedia* | $1.44 \times 10^{-1}$ |
| Iceland | *Bacillus soli* | $5.57 \times 10^{-1}$ |
| Iceland | *Microbacterium arborescens* | $6.85 \times 10^{-1}$ |
| Iceland | *Prolinborus fasciculus* | $1.86 \times 10^{-1}$ |
| Silwood | *Cedecea lapagei* | $1.54 \times 10^{-2}$ * |
| Silwood | *Flavobacterium tyrosinilyticum* | $3.40 \times 10^{-2}$ * |
| Silwood | *Scandinavium goeteborgense* | $2.62 \times 10^{-2}$ * |
| Silwood | *Raoultella planticola* | $7.16 \times 10^{-2}$ |
| Silwood | *Serratia liquefaciens* | $7.52 \times 10^{-2}$ |
| Silwood | *Pedobacter trunci* | $3.00 \times 10^{-1}$ |
| Silwood | *Psychrobacillus glaciei* | $1.42 \times 10^{-1}$ |
| Silwood | *Pseudomonas fragi* | $3.44 \times 10^{-1}$ |
| Silwood | *Sphingobacterium kitahiroshimense* | $1.36 \times 10^{-1}$ |
| Silwood | *Microbacterium algeriense* | $3.49 \times 10^{-1}$ |
| Silwood | *Microbacterium natoriense* | $9.30 \times 10^{-2}$ |
| Silwood | *Paeniglutamicibacter antarcticus* | $7.32 \times 10^{-2}$ |
| Silwood | *Rhodococcus wratislaviensis* | $1.53 \times 10^{-1}$ |
| Silwood | *Priestia flexa* | $2.38 \times 10^{-1}$ |

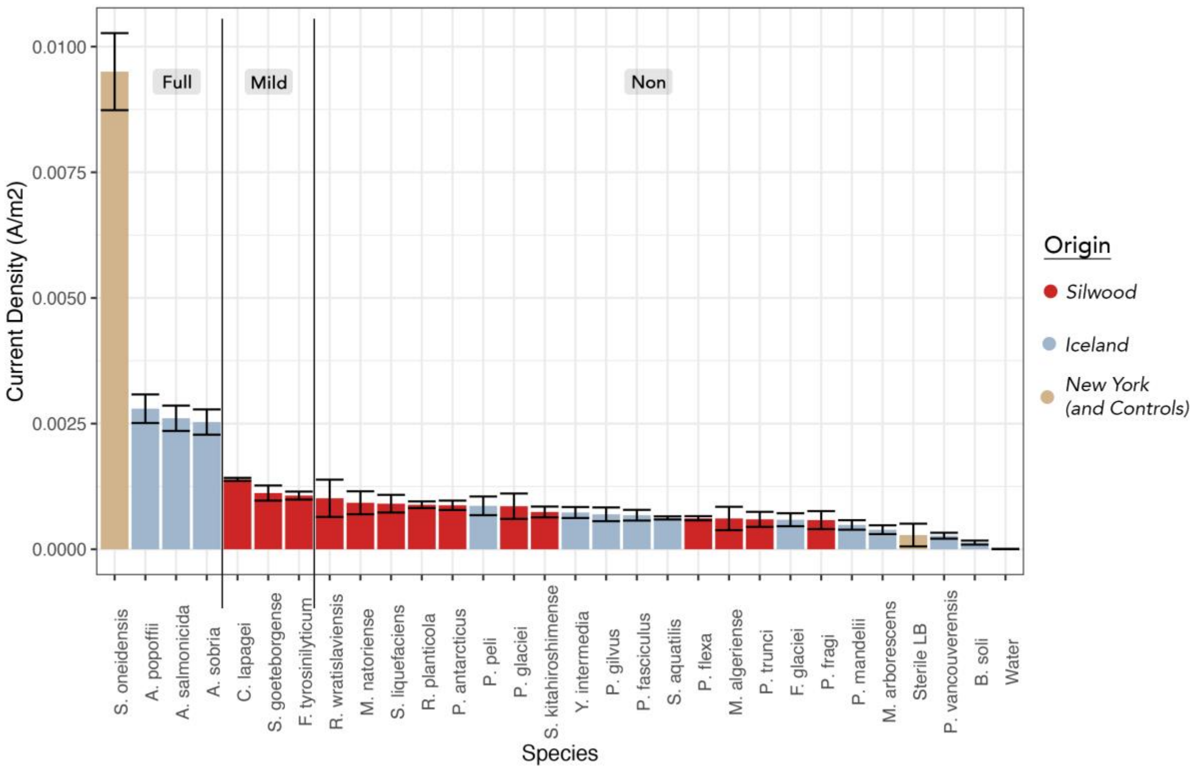


**Figure 3:** The electrical performance for each of the monocultured isolate with their exoelectrogenic status and origin assigned. Current density was calculated using Ohm's law and electrode surface area measurements. Each replicate for each inoculant was biologically independent, typically being performed in duplicate experiments. A minimum of four biological replicates was performed for each isolate. For each independent biological replicate, the median current density was determined. The displayed data is the mean of these medians with the error bars displaying the standard error of the mean.

#### Coulombic Efficiency

A common productivity metric for fuel cells is coulombic efficiency which describes the quantity of electrons that successfully make it to the anode and thus a circuit [9]. The coulombic efficiency of each assessed isolate was determined, as per equation (2), originating from Oh, Min & Logan (2004) [10]. A constraint with this measurement is the non-systematic methods of calculation, further highlighting the absence of standardised MFC inoculant performance measure [10,11]. Additionally, such calculations require the molarity of minimal medias (eg: acetate) which is challenging to determine for LB and its constituents [10].

Figure 4 shows very low coulombic efficiencies for the assessed isolates, with the highest being exhibited by *S. oneidensis* MR-1 at approximately 0.07%. A t-test was used to compare the coulombic efficiency of each isolate to the sterile LB control, with *S. oneidensis* MR-1 being the only isolate to yield a significant result ($p$ = 0.002). While some studies have achieved a coulombic efficiency of 97%, other studies have achieved 0.04%, highlighting the varying influence that taxa, operating conditions and substrate have on coulombic efficiency [9,12–14].

$$E = \frac{C_{EX}}{C_{TH}} \; x \; 100 \qquad (2)$$

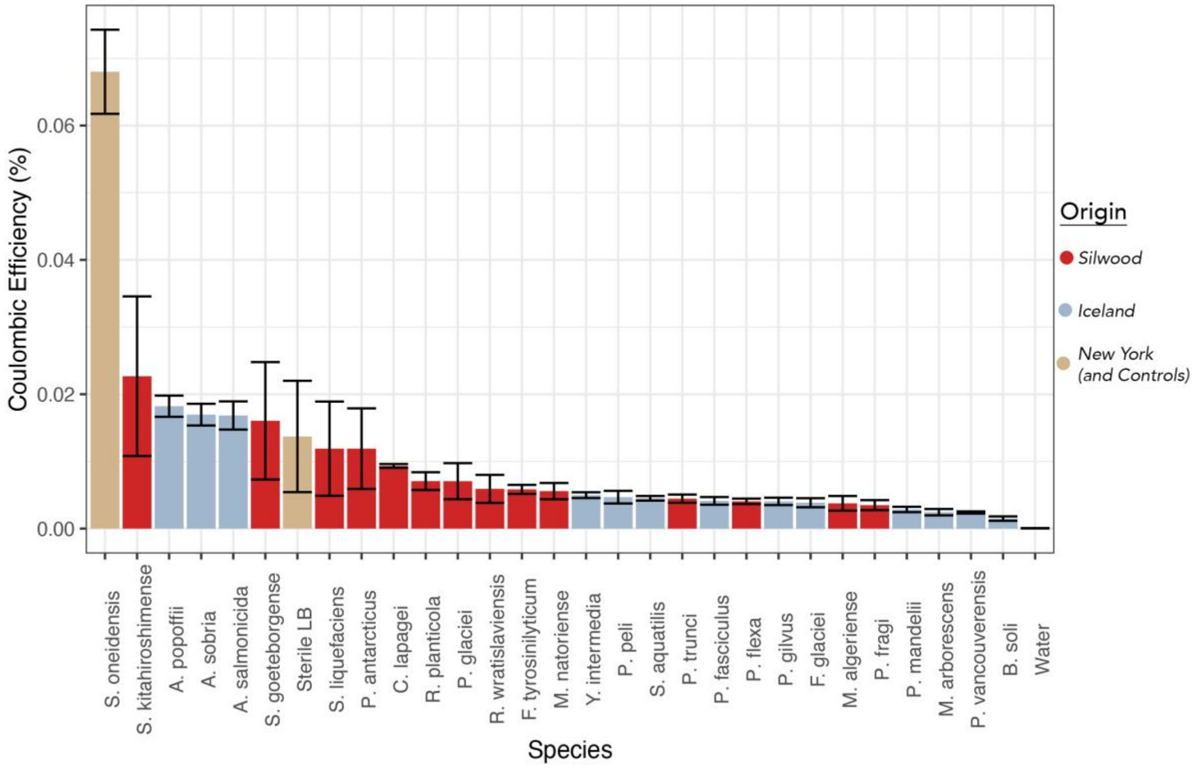


**Figure 4:** The coulombic efficiency, as a percentage, of each of the monocultured isolates. Efficiencies were calculated using the equation 3. Each replicate for each inoculant was biologically independent, typically being performed in duplicate experiments. A minimum of four biological replicates was performed for each isolate. For each independent biological replicate, the median coulombic efficiency was determined. The displayed data is the mean of these medians with the error bars displaying the standard error of the mean. Further species information is displayed in Table 1.
